# The gateway into Remote Oceania: new insights from genome-wide data

**DOI:** 10.1101/190843

**Authors:** Irina Pugach, Ana T. Duggan, D. Andrew Merriwether, Françoise R. Friedlaender, Jonathan S. Friedlaender, Mark Stoneking

## Abstract

A widely accepted two-wave scenario of human settlement of Oceania involves the first out-of-Africa migration ca 50,000 ya, and one of the most geographically-widespread dispersals of people, known as the Austronesian expansion, which reached the Bismarck Archipelago by about 3,450 ya. While earlier genetic studies provided evidence for extensive sex-biased admixture between the incoming and the indigenous populations, some archaeological, linguistic and genetic evidence indicates a more complicated picture of settlement. To study regional variation in Oceania in more detail, we have compiled a genome-wide dataset of 823 individuals from 72 populations (including 50 populations from Oceania) and over 620,000 autosomal SNPs. We show that the initial dispersal of people from the Bismarck Archipelago into Remote Oceania occurred in a “leapfrog” fashion, completely by-passing the main chain of the Solomon Islands, and that the colonization of the Solomon Islands proceeded in a bi-directional manner. Our results also support a divergence between western and eastern Solomons, in agreement with the sharp linguistic divide known as the Tryon-Hackman line. We also report substantial post-Austronesian gene flow across the Solomons. In particular, Santa Cruz (in Remote Oceania) exhibits extraordinarily high levels of Papuan ancestry that cannot be explained by a simple bottleneck/founder event scenario. Finally, we use simulations to show that discrepancies between different methods for dating admixture likely reflect different sensitivities of the methods to multiple admixture events from the same (or similar) sources. Overall, this study points to the importance of fine-scale sampling to understand the complexities of human population history.

## INTRODUCTION

The Pacific is a vast region, encompassing an entire hemisphere of our planet, and the human settlement of the far-flung Pacific Islands has long been of intense interest. A convenient division of the Pacific consists of Near and Remote Oceania, with the border located between Makira, the most easterly island of the main Solomon Island Archipelago, and the islands of Santa Cruz (politically part of the Solomon Islands), Vanuatu, and New Caledonia (Fig.S1). Fossil, archaeological, and genetic evidence indicate that Near Oceania was initially colonized by 45,000 – 50,000 years ago (Groube et al. 1986; Roberts et al. 1990; O’Connell & Allen 2015; Malaspinas et al. 2016) or possibly even earlier (Clarkson et al. 2017); at this time, sea levels were considerably lower and most of Near Oceania (Australia, New Guinea, and many of the nearby islands) were connected as a single landmass called Sahul. However, Sahul was never connected to the continental Asia landmass (Sunda), which at the time encompassed the present islands of Sumatra, Java, and Borneo, and so humans had to cross water in order to colonize Sahul. These crossings would have been mostly intervisible, meaning that there would have been some indication of land ahead before losing sight of the land behind, and so may not have required particularly sophisticated boating technology or navigational skills. By contrast, the initial colonization of Remote Oceania would have involved crossing at least 400 km of open ocean, and was only accomplished about 3,500 years ago (Spriggs 2003); colonization of the other islands of Remote Oceania involved crossing thousands of kilometers of open ocean.

The languages spoken today in Near Oceania are classified as either Papuan or Austronesian. The indigenous Papuan languages are so deeply-rooted and varied that the relationships between them are still poorly understood (Hunley et al. 2007; Pawley 2007); they do not constitute a language family in the usual sense that the languages can be demonstrably related and they are also quite distinct from the Austronesian languages. By contrast, Austronesian languages can be demonstrably related (Blust 1999; Gray et al. 2009). Their introduction to Oceania was part of an expansion that probably started from Taiwan about 5,000 years ago (Bellwood 2004; Ko et al. 2014) and arrived in the Bismarck Archipelago about 3,400 years ago, in association with the appearance of Lapita pottery (Spriggs 2003; Bellwood & Dizon 2005). This Austronesian expansion then continued to, and was the first to cross, the border to Remote Oceania, arriving there by 3,200 years ago (Kirch 2000; Bellwood 2004).

While there is growing consensus for the above “dual-wave” model for the colonization of Near and Remote Oceania, many questions remain. For example, there is an approximately 40,000-45,000 year hiatus and regional isolation between the initial colonization of Near Oceania and the Austronesian expansion; are there any detectable signals in the genomes of modern Oceanian populations of any other migrations during this time period? While some studies of mtDNA variation have claimed to find evidence for pre-Austronesian contact between Southeast Asia and Near Oceania (Soares et al. 2011), the most detailed study to date of complete mtDNA genome sequences from Oceania estimated that mtDNA sequences attributable to any such pre-Austronesian contact constitute at most 2% of the current Oceanian mtDNA gene pool (Duggan et al. 2014). Y chromosome studies conducted to date in Near Oceania lack sufficient resolution to address the issue of pre-Austronesian migrations (Kayser et al. 2000; Kayser et al. 2006; Delfin et al. 2012).

Another question concerns the timing of the contact that occurred between the Austronesian-speaking groups and the resident populations of Near Oceania (referred to here collectively as Papuans, while not implying any cultural, linguistic, or biological unity of these populations), prior to the colonization of Remote Oceania. The contemporary populations of Remote Oceania all carry both Asian-related and Papuan-related ancestry (Kayser et al. 2000; Kayser et al. 2006; Kayser et al. 2008; Friedlaender et al. 2008; Wollstein et al. 2010; Skoglund et al. 2016), and various methods for dating this signal of admixture provide estimates of about 3,500 years ago (Wollstein et al. 2010; Pugach et al. 2011). This suggests that admixture occurred shortly after the arrival of Austronesian speakers in Near Oceania (most likely in the Bismarck Archipelago) and prior to the colonization of Remote Oceania. However, a recent study of ancient DNA obtained from skeletons dating to the initial occupation of Vanuatu and Tonga did not find any evidence of Papuan ancestry, and moreover dated the admixture (using the ALDER method) in contemporary Tongans to only about 1,500 – 2,300 years ago (Skoglund et al. 2016). These results suggest that the initial colonization of Remote Oceania was by people whose ancestors had not yet mixed with Papuans, and that Papuan ancestry was introduced to Remote Oceania by more recent migrations. Indeed, it had previously been observed that there is higher and more variable amounts of Papuan ancestry in Fiji than elsewhere in Remote Oceania, which can be explained by additional migration(s) of Papuan people that reached as far as Fiji (Wollstein et al. 2010). Nonetheless, there is still a conflict between the dates of the Papuan-Asian admixture signal in current Polynesians of 1,500-2,300 years ago estimated by the ALDER method (Skoglund et al. 2016) vs. dates of ~3,500 years ago estimated by other methods (Wollstein et al. 2010; Pugach et al. 2011).

A further question is whether the movement of people from Near to Remote Oceania followed a simple wave-of-advance model (i.e., proceeding from New Guinea to Santa Cruz/ Vanuatu/ New Caledonia via the main Solomon Islands archipelago, and then from this region to the rest of Remote Oceania), or whether movement was more complicated. There are already some indications that migration was more complicated than a simple wave-of-advance model; for example, obsidian from the Bismarck Islands is found in archaeological sites on Santa Cruz but not elsewhere in the Solomon Islands (Sheppard & Walter 2006), suggesting contact between Santa Cruz and the Bismarcks that bypassed the main Solomon Island archipelago. Santa Cruz is also unusual in having a much higher frequency of Papuan mtDNA and Y chromosome haplogroups than anywhere else in Remote Oceania (Friedlaender et al. 2002; Delfin et al. 2012; Duggan et al. 2014), and the diversity of Papuan mtDNA sequences is too high to be explained by a simple bottleneck or founder event (Duggan et al. 2014). Instead, it appears as if there was substantial contact between Santa Cruz and Near Oceania that did not penetrate much further into Remote Oceania.

To address these and other issues concerning the relationships among and between populations of Near and Remote Oceania, we have assembled a dataset of 823 individuals from 72 populations, including 50 populations from Near and Remote Oceania (Table.S1), all genotyped for ~620,000 SNPs on the Affymetrix Human Origins Array (Patterson et al. 2012). Our analyses indicate substantial complexity and waves of contact during the movement of people through Near Oceania to Remote Oceania. We also use computer simulations to address the discrepancy in dates for the Papuan/Asian admixture signal obtained by ALDER vs. other methods, and show that this is likely to reflect the different effect of multiple episodes of admixture from the same (or similar) source populations on these methods. Overall, our study highlights how combining sampling on a fine scale with dense genome-wide data can illuminate new aspects of the peopling of the Pacific.

## RESULTS AND DISCUSSION

### Large-scale population structure

First, to obtain an overview of the data structure and potential associations between the samples in the dataset, we started with principal components analysis (PCA) (as implemented in the StepPCO software (Pugach et al. 2011)). PCA performed on the entire dataset (Fig.S2) reveals that the first principal axis (PC) is driven by differences between the populations of Eurasia and the highlands of Papua New Guinea (NGH), while the second axis separates Europe and Asia. Oceanian populations form a cline from NGH to East Asia (notably, the Taiwan aboriginals), with the westernmost populations of Near Oceania falling closer to NGH, and Polynesian populations falling closer to East Asia (Fig.S2). More details become apparent when we limit the analysis to populations from Oceania and Southeast Asia (Fig.1). In this analysis, while the first PC captures the variation between populations from Oceania and Asia, the second is driven by the differences between the NGH and Bougainville, with populations from the Bismarcks and Australia falling in the middle. The Polynesian Outliers (PO) fall close to the Tongans, and are placed in between Bougainville and SE Asia. Although it has been previously shown that when applied to human genetic data from some geographic regions, such as Europe, the first two principal axes correlate closely with geography (Lao et al. 2008; Novembre et al. 2008), we do not observe such correlation for populations of the Solomon Islands (SI). In particular, while the populations from the western SI fall close to Bougainville (politically part of PNG, but geographically the largest and westernmost island in the SI chain), the populations from the eastern part of the chain are markedly shifted towards the Bismarcks Archipelago (BA,which is geographically located northwest of Bougainville; Fig. 1 and Fig.S1), with the easternmost islands of the SI chain falling closest to the BA populations. Strikingly, the samples from Santa Cruz (politically part of the Solomon Islands, but geographically separate from them, in Remote Oceania) fall directly adjacent to populations from island New Britain (the largest island in the BA), and this overlap is not coincidental, as it continues to be seen in higher PCs (Fig.S3). This result of eastern SI falling closer in PC space to the BA persists if the analysis is carried out on the subset of the data which includes only the Oceanian populations (Fig.S4). Furthermore, as is seen in Fig.S4, in this analysis Santa Cruz completely overlaps with the BA samples along the first PC, but shows a slight shift towards NGH along the second PC.

**Fig.1.**
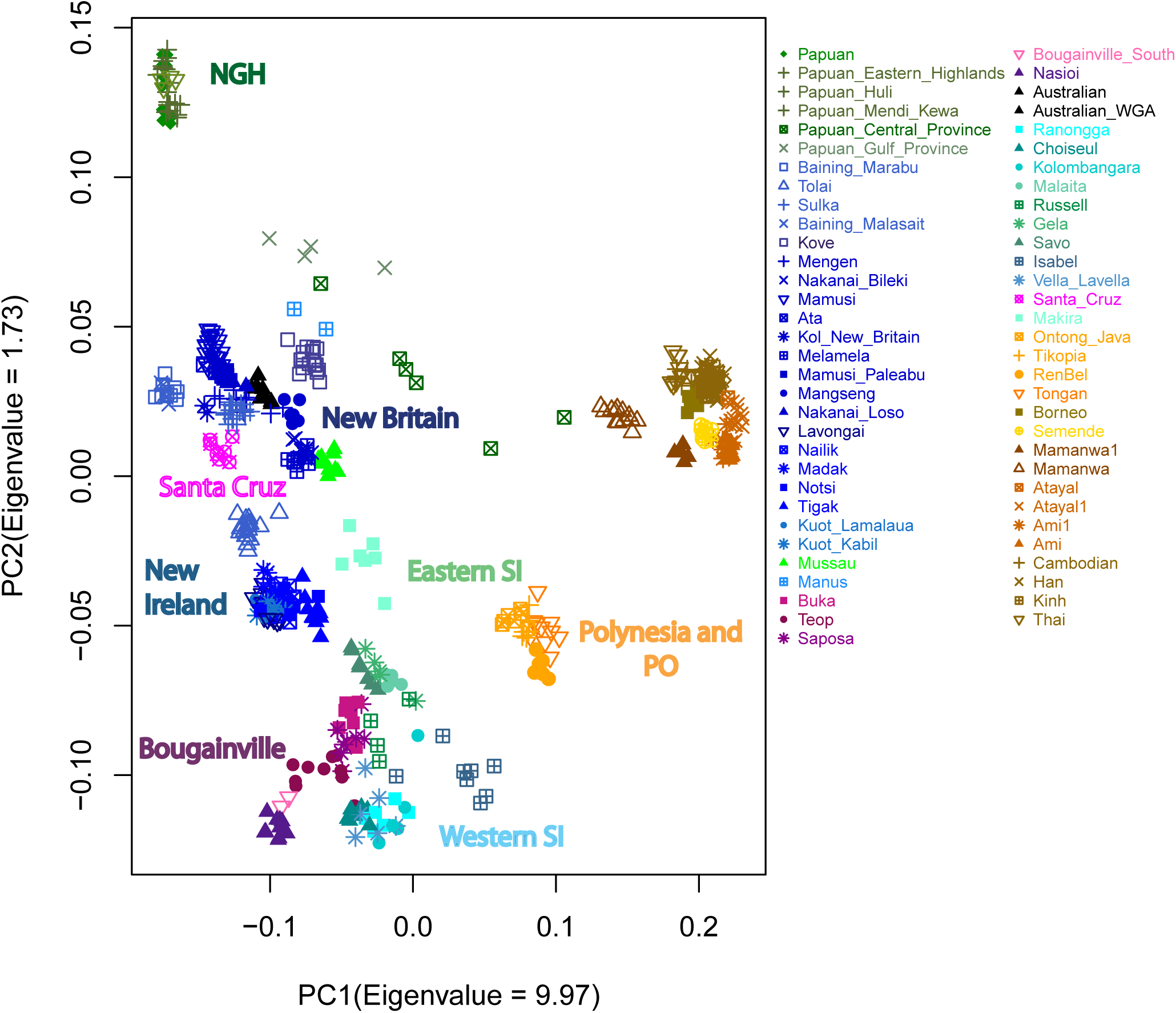
Results of the PC analysis showing the genetic structure captured by the first two principal components. Each colored label represents an individual.

Next, to further examine broad genetic structure based on haplotypes and not individual SNPs, we used BEAGLE (Browning & Browning 2007; Browning & Browning 2011) to phase the data and infer segments inherited by each pair of individuals from the most recent common ancestor that are identical by descent (i.e. without recombination). We then performed PC analysis on the inverse of the similarity matrix calculated from the number of segments shared identical by descent (IBD) by each pair of individuals (Pugach et al. 2016). Since the detection of IBD segments based on genotype data, rather than sequence data, is biased towards detection of longer fragments (Ralph & Coop 2013) we expect this analysis to reveal the history of recent interactions (occurring mostly in the past 4,000 years) (Ralph & Coop 2013). The results (Fig.3)indicate that at this time-scale we lose all differentiation between populations of the Solomon Islands, Bougainville, Polynesian Outliers, and some populations from the BA (namely New Ireland), which indicates substantial haplotype sharing and hence would argue for considerable recent gene flow between these locations. Notably, however, this signal of close recent relatedness is not observed for samples from Santa Cruz, which instead continue to exhibit closer distances to populations from New Britain (BA) and NGH (Fig.3).

**Fig.2.**
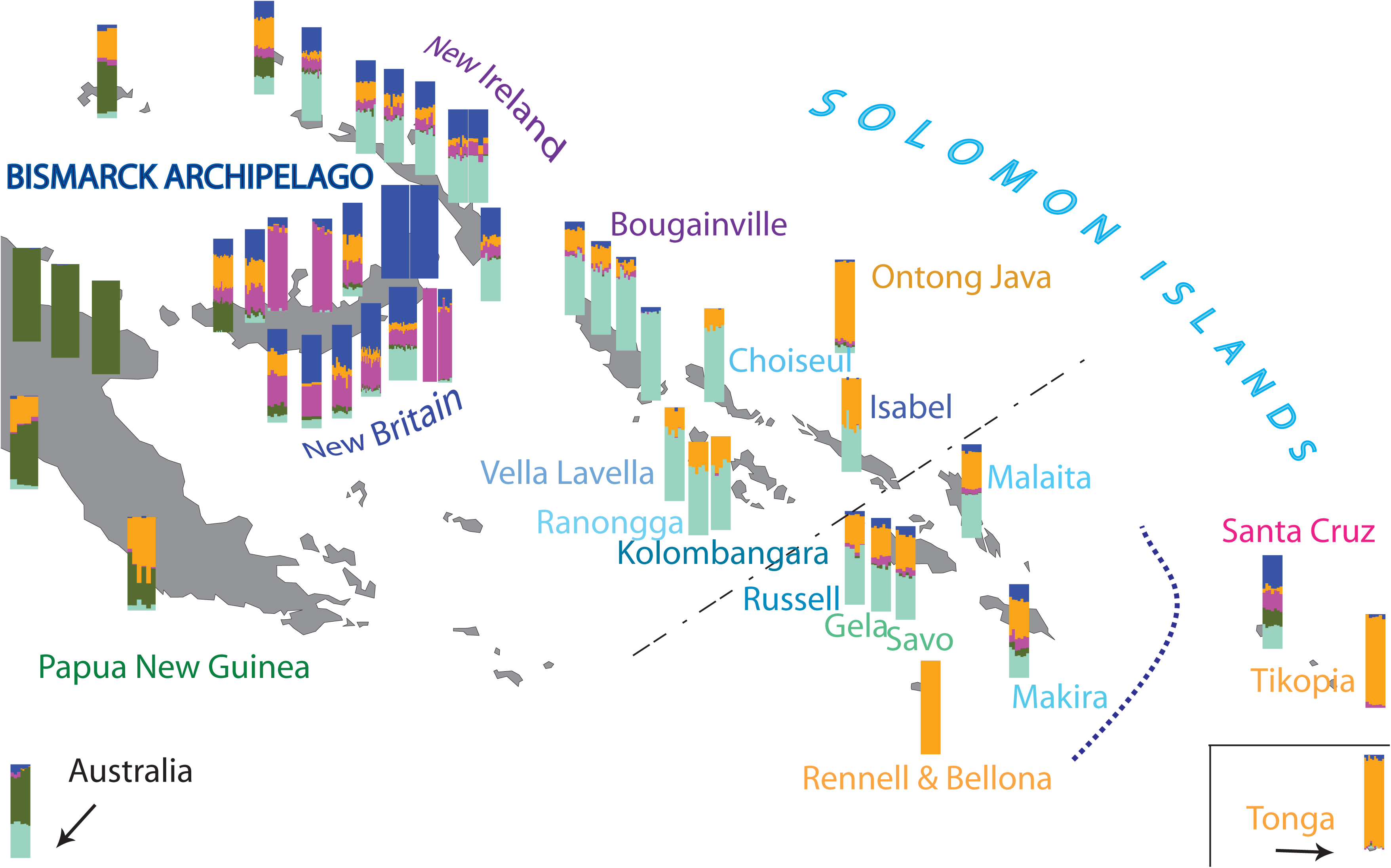
DMIXTURE results for K = 5 showing the approximate location of the Oceanian populations included in this study. For reasons of space the location of aboriginal Australians and Tongans does not correspond to their true location (which can be seen in Fig.S1). The curved dotted line marks the biogeographic boundary between Near and Remote Oceania, while the straight broken line denotes the Tryon-Hackman linguistic divide.

**Fig.3.**
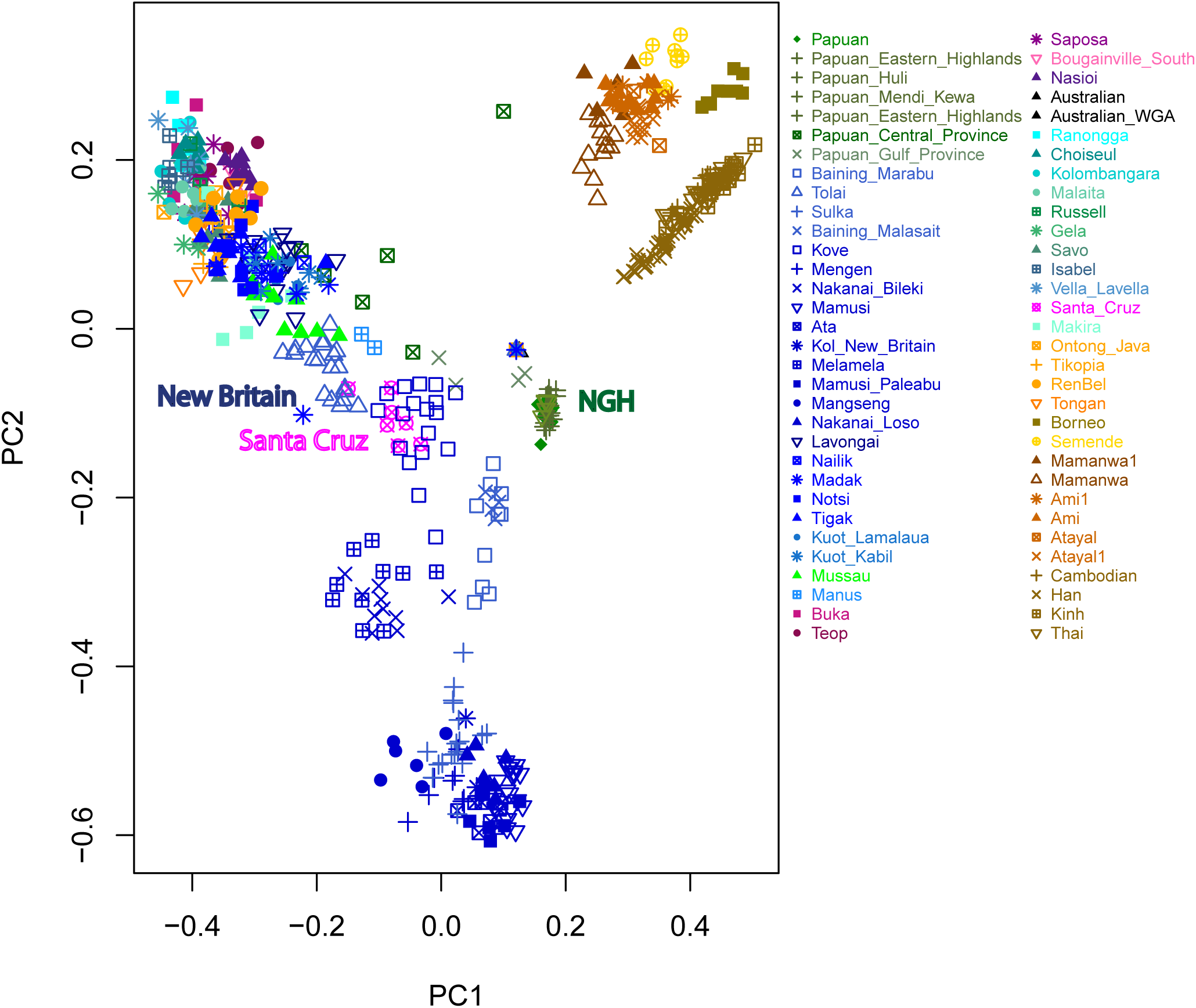
Results of the PC analysis showing recent relatedness based on the number of IBD blocks shared between individuals.

To further investigate the general patterns of relatedness between populations in the dataset, we used a clustering approach, ADMIXTURE (Alexander et al. 2009), which models each individual's genome as having originated from a fixed number of hypothetical ancestral populations. When running the analysis on the complete dataset, we explored K = 3 through K=12 (Fig.S5A), with the lowest cross-validation error (and hence the best predictive value for K) observed for K=10 (Fig.S6A). At this value of K, five of the ancestry components can be roughly ascribed to Europe, south Siberia, India, Onge and Asia, with the remaining five ancestry components belonging to Oceania (Fig.S7). As the Human Origins Array contains 13 separate ascertainment panels, we also performed the analysis for each separate panel for K = 10 (Fig. S8). In general, higher levels of Papuan ancestry (dark green component) are inferred across the Bismarcks and eastern Solomons than is inferred for these populations with the full set of markers, which is probably due to the limited resolution offered by only a subset of SNPs to differentiate between the closely related Papuan-like ancestries (blue, aqua and pink components). Interestingly, with Papuan ascertainment panels there is 15-20% less Papuan and 5% more Indian ancestry inferred in Australia, which is supportive of previous claims of Indian gene flow to Australia (Pugach et al. 2013).

Since the analyses of the full set of SNPS as well as the individual ascertainment panels reveal little impact of Eurasian populations on genetic structure in Oceania, we repeated the ADMIXTURE analysis for the Oceanian data only (Fig.S5B). For this subset the lowest cross-validation error is seen for K=5 (Fig.S6B). Similar to the results obtained for the full dataset (Fig.S7), the five Oceanian components could broadly be assigned to: New Guinea (dark green; seen at highest frequency in NGH): Polynesia (orange; seen at highest frequency in Polynesian outliers and Tonga), Bougainville (aqua; seen at highest frequency in Nasioi), and two components found at highest frequency in the BA, namely in the Baining (blue) and Mamusi (pink) populations of New Britain (Fig.S9).

When the results of the ADMIXTURE analysis for each population are superimposed onto a geographic map (Fig.2), several patterns become apparent. Firstly, the Papuan ancestry component (dark green) is found at high or appreciable frequency only in the west - on the main island of New Guinea, in Australia, and in some coastal populations of New Britain; it is almost completely absent from New Ireland and does not appear at all across the SI, with the exception of the easternmost islands of the chain - Makira and Santa Cruz - where it appears at substantial frequency. The two components which appear at high frequency on New Britain (blue and pink) are always observed together; they are prevalent throughout the BA and are almost absent from the western SI, yet they reach a substantial frequency in the eastern SI, accounting for more than 50% of the inferred ancestry in Santa Cruz, ~30% in Makira, and ~20% in other populations of the eastern SI. The aqua component, which is present at high frequency on Bougainville, is ubiquitous east of mainland New Guinea and seems to exhibit a clinal pattern with its frequency decreasing with increasing distance from Bougainville. The Polynesian component (orange) also seems to exhibit a geographical gradient, with its frequency diminishing in an east to west direction. Some small islands (e.g. Manus, Mussau) and coastal populations (e.g. Kove, Melamela) from western New Britain do not follow this cline, having a third of their ancestry assigned to this Polynesian component. An exception at the opposite extreme are individuals from Santa Cruz, who have very little (on average 5%) of this component. In addition, the aboriginal Australian samples reveal a baffling signal of admixture, showing both Papuan and Bougainville ancestry components in roughly equal proportion; the interpretation of this signal is considered below.

Taken together these results tentatively suggest different historical trajectories for western vs eastern SI (Fig.1, Fig.2); confirm the population of Santa Cruz as a genetic outlier with respect to its neighbors (Fig.1, Fig.2), as found in previous studies of uniparental markers (Friedlaender et al. 2002; Delfin et al. 2012; Duggan et al. 2014); and suggest both long-term isolation for some populations (e.g. Baining, Mamusi, Nasioi and NGH) and ubiquitous recent post-Austronesian gene flow for others (SI, except Santa Cruz and the PO, Fig.3). However, because both PCA and ADMIXTURE are descriptive analyses that do not directly inform about the underlying historical processes, in the subsequent sections we apply additional methods to first validate and then to expand these tentative findings.

### Evaluating diverse signals of admixture in Oceania

Despite the misleading name of the ADMIXTURE software and the usual interpretation given to the results of this and other similar clustering approaches (e.g. STRUCTURE and *frappe*), assignment of an individual's genome to two or more ancestry components inferred by the analysis suggests, but does not necessarily demonstrate, admixture. Furthermore, other demographic processes not involving admixture, such as drift (Hudjashov et al. 2017), recent bottlenecks (Falush et al. 2016) as well as uneven sampling (Puechmaille 2016) could produce ADMIXTURE results indistinguishable from those which reflect actual admixture. To formally evaluate signals of admixture in Oceanian populations we first applied the 3-population test (Patterson et al. 2012), in the form of f3(C; A, B), which tests for “treeness”, and where a significantly negative value of the f3 statistic would imply that population C has descended from an admixture event between A and B. We find signals of admixture for most, but not all, Oceanian populations (Table S2). The most significantly negative f3 values are observed when the admixture event involves NGH, Baining or Nasioi populations (all of which are assigned their own ancestry component in the ADMIXTURE analysis) and Taiwanese aboriginals, probably reflecting Austronesian expansion (Spriggs 2003). Out of the source populations which exclude Asia, the most significant results are obtained with either Tonga or Isabel (western SI) as one of the sources (except for Papuans and Saposa, where the most significant result is obtained with Sulka and Ontong Java, respectively). Since this result is confounded by the fact that both Tongans and Isabel show evidence of admixture from Taiwan, it remains to be further evaluated whether or not this signal reflects additional gene flow. In Table.S2 we report for each population the top three results with the highest significance score, followed by the top three results when the source populations do not include Asia. Importantly, admixture was not inferred via the f3 statistic for any of the populations which have been ascribed their own component in the ADMIXTURE analysis at K=5 (namely NGH, Baining and Mamusi groups, Nasioi, Tikopia and Rennell and Bellona).

While significantly negative f3 scores are taken as clear evidence of admixture, nonsignificant tests do not necessarily indicate the absence of admixture, since processes such as genetic drift following admixture could mask the signal entirely (Patterson et al. 2012). Similarly, in analyses such as ADMIXTURE, recent genetic drift could make admixed drifted populations look distinct and hence assigned pure ancestry components, while the unadmixed sister populations might by contrast appear admixed, simply because they have a larger population size and greater inter-individual genetic variation (Falush et al. 2016). This is probably the case for the aboriginal Australians in this dataset, who contrary to numerous previous studies (e.g McEvoy et al. 2010; Pugach et al. 2013; Malaspinas et al. 2016) appear to have admixed NGH and Bougainville ancestry (Fig. 2). However, none of the f3 scores are significant for the aboriginal Australians, and so the most likely explanation is that Australia, NGH, and Bougainville all share ancestry but NGH and Bougainville experienced more genetic drift after population divergence. This enhanced drift is more easily detected as a separate ancestry component in the ADMIXTURE analysis, and since aboriginal Australians share ancestry with both ancestry components, they are (mistakenly) assigned as admixed.

Next, we used TreeMix (Pickrell & Pritchard 2012) to build a population tree and identify admixture events (which in this approach are added sequentially to the constructed tree until the residuals between the observed data and the fitted data are reduced). First, we performed the TreeMix analysis on the full dataset of 75 populations, adding the Yorubans as an outgroup. The inferred maximum-likelihood tree largely groups the populations as expected (Fig.S10A), but with Santa Cruz grouping with populations from New Britain, and not the Solomon Islands. Positive residuals of the standard error capture pairs of populations that are more closely related to each other than is suggested by the inferred tree, and are thus candidates for gene flow (Pickrell & Pritchard 2012). The residuals (Fig.S10B) show the greatest error for: a) the Australian Aboriginals vs Indian and European populations: b) the Australian Aboriginals vs the Papuan Highlanders; c) Papuan Highlanders vs coastal populations of New Guinea, Kove from the coast of northwest New Britain and Manus - the largest of the Admiralty Islands to the north of PNG; d) western SI vs Bougainville; e) Taiwanese Aboriginals, Mamanwa (Philippines), Semende (Sumatra) and to a lesser extent island southeast Aisa in general vs Polynesians and Isabel; and f) Papuan Gulf Province vs Europeans. However, when we start adding migration edges to the tree the results appear to be meaningless (Fig.S11A). This is most likely explained by the large number of populations that TreeMix is trying to fit simultaneously, which is prohibitive for disentanglement of complex admixture histories (Lipson et al. 2013). Indeed, when we reduce the dataset by omitting the populations from the BA, the migration edges inferred by TreeMix recover all the purported signals of admixture, such as gene flow from the Taiwanese Aboriginals to Polynesians, from NGH to the coastal groups and to the eastern SI, and from Bougainville to the western SI (Fig.S11B).

### Sharing of IBD segments

To more clearly elucidate the patterns of recent relatedness, local migrations and recent genetic drift, we analyzed IBD blocks, i.e. segments of DNA that were inherited without recombination by each pair of individuals from their most recent common ancestor (Powell et al. 2010; Ralph & Coop 2013). We expect that recent gene flow between two populations would result in them sharing a higher number of longer IBD segments in comparison to others. Also, because the genomes of individuals in a drifted population would exhibit a congruence of genealogies, we expect to see more IBD segments shared by individuals within such populations (Gusev et al. 2012) as opposed to individuals from a population which did not experience recent drift (Ralph & Coop 2013; Harris & Nielsen 2013).

When we consider the sharing of IBD segments within each population, we observe that the majority of populations assigned their own ancestry component in the ADMIXTURE analysis also exhibit extensive IBD-segment sharing within the population (Fig.4). This pattern is further corroborated by the genome-wide analysis of runs of homozygosity (ROH), which are uninterrupted chromosome regions in which all loci in an individual are homozygous, due to identical chromosomal segments being inherited from both parents. Thus, as with IBD segments, ROHs provide a useful measure of parental relatedness and historical population isolation (e.g. McQuillan et al. 2008). Again, all populations in the dataset that were assigned their own ancestry component in the ADMIXTURE analysis also exhibit an excess of very long ROHs in comparison with other populations (Fig.S12), which is typical of isolated populations that underwent a strong bottleneck (Kirin et al. 2010).

**Fig.4.**
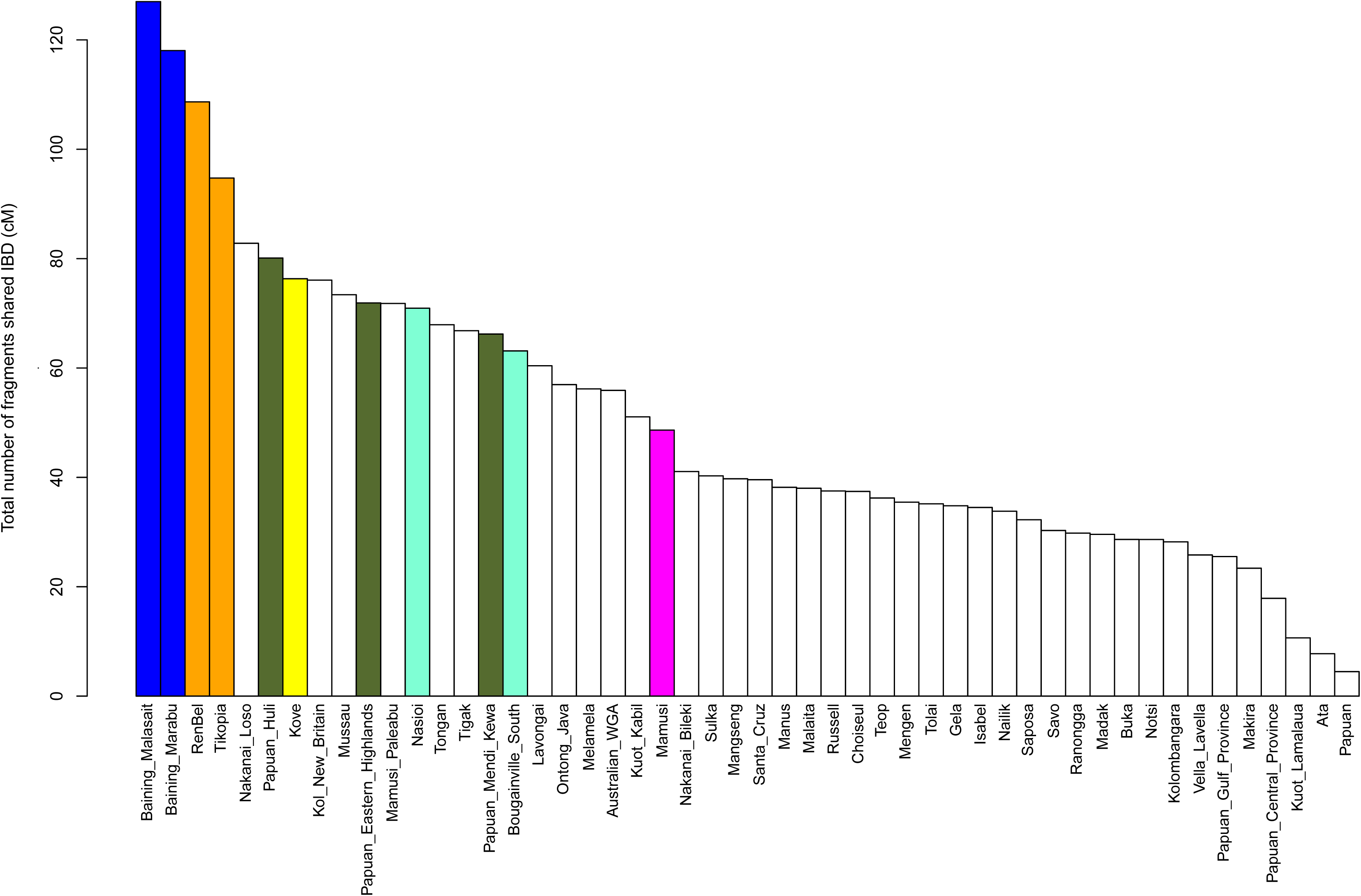
Number of IBD blocks shared within populations. The blue, orange, dark green, aqua and pink colors identify the five main components inferred in the ADMIXTURE analysis at K=5. The colored bars indicate populations which have their own ancestry component assigned to them in at K=5 (yellow corresponds to the sixth main component, which at K=6 is assigned to Kove (see Fig.S5B)

In terms of between population IBD sharing, we observe that most populations of New Britain (the largest island in the BA) show substantial sharing of IBD segments with their neighbors, namely populations from New Ireland (BA), Bougainville, and the eastern part of the SI chain, and also with the PO and Tongans. The exceptions are: a) the Baining groups, who share only with the neighboring Sulka and Tolai, and with Santa Cruz (one of the Baining groups also shows sharing with Polynesians); and b) the Nakanai Loso, Ata and the Mamusi groups, who share only with their neighbors, and in agreement with the ADMIXTURE result (Fig.2) reveal no connection to New Ireland, the SI or Polynesia (Fig.S13A). In general, all populations from across New Ireland share IBD segments extensively with their geographic neighbors (namely the Kuot groups, Nailik, Notsi and Madak, which are located more centrally than the Tigak, who come from the northwestern tip of the island), as well as with Sulka, Tolai and Kove from New Britain. The length of the IBD segments reflects that the contact must have been very recent. In contrast the length of the IBD segments shared with the SI and with Tonga and the PO suggests older ties; except for the Tigak who reveal a more recent sharing with Isabel (SI) and with Tongans. Like the coastal Tigak population, the Lavongai people from New Hanover, (located a few kilometers northwest of New Ireland) extensively share IBD segments with populations from New Ireland and New Britain; they also reveal a more recent contact with Bougainville and the western SI, and with the Polynesians (Fig.S13B). In contrast to populations from New Ireland, but similarly to Lavongai, the two populations residing on Manus and Mussau (the islands located furthest to the north) not only share IBD segments with the other populations from the BA, the Solomon Islands, PO and the Tongans, but also with the Papuans (consistent with results of the ADMIXTURE analysis, (Fig.2)). However, the length of shared IBDs is much lower in Manus, who share long IBDs only with the Nakanai Bileki from New Britain, with Isabel (SI) and with Tongans (Fig.S13C). Bougainville populations (Nasioi, Buka, Saposa and Teop) show substantial sharing with each other and across the western SI, with little/moderate sharing with populations from eastern SI. Except for the Teop, they all share recent connections with the Tolai (New Britain) and Lavongai. Interestingly, Nasioi exhibits an equal amount of sharing with all populations of Polynesian ancestry, while the others share much longer tracts with the Tongans and Ontong Java. Similarly, the islands in the western SI (Ranongga, Choiseul, Kolombangara and Vella Lavella; Isabel is discussed separately below) show substantial sharing with populations from Bougainville and with each other, but much less sharing with populations from the eastern SI. In addition, Choiseul and Kolombangara show recent links with the Tolai (New Britain). The western SI populations share more with Tonga than with the PO. By contrast, populations from the eastern SI (Russell, Gela, Savo, Malaita and Makira) show more Papuan-related ancestry than the western SI and also exhibit more sharing with the BA than with Bougainville or the western SI (except Isabel, which is discussed below). They also do not show any particular recent links to Santa Cruz. In addition, like many other islands across the SI chain, Makira and Savo show a recent genetic relationship with the Tolai from New Britain (Fig.S13D).

Isabel and Santa Cruz depart from these general patterns of ancestry sharing for the SI. Isabel shows extensive recent genetic sharing with most populations from the BA and the SI (Santa Cruz being the notable exception), and with all sampled populations of Polynesian ancestry. Santa Cruz shares more segments IBD with New Britain than with New Ireland, Bougainville, or eastern or western SI (including Isabel).

Peculiarly, Santa Cruz exhibits recent genetic ties with the Tolai, as do some other islands in the Solomons. Santa Cruz also shows equally low levels of sharing with all populations of Polynesian ancestry; this nonspecific sharing may indicate that Santa Cruz did not experience any additional post-Austronesian gene flow; by contrast, other populations exhibit ubiquitous sharing of long IBD segments with either Tongans (e.g. Sulka, Tigak, Choiseul, Savo), or Tongans and the PO (e.g. Buka, Saposa, Isabel, Makira), suggestive of contact post-dating the Austronesian expansion (Fig.S13E).

In summary, the analyses based on the number and length of segments shared IBD highlight a) a considerable recent post-Austronesian expansion gene flow, which is diverse both in terms of its sources and direction, across many of the islands in Near Oceania; b) different patterns of sharing in western vs eastern Solomons (Fig.5), characterized by substantial local sharing of long IBD segments in the western Solomons vs fairly restricted local sharing in the eastern Solomons; c) special roles of Isabel islanders and the Tolai people of the Gazelle Peninsula, New Britain, who show recent genetic ties with many populations across Oceania; d) notably low levels of sharing between Santa Cruz and other populations.

**Fig.5.**
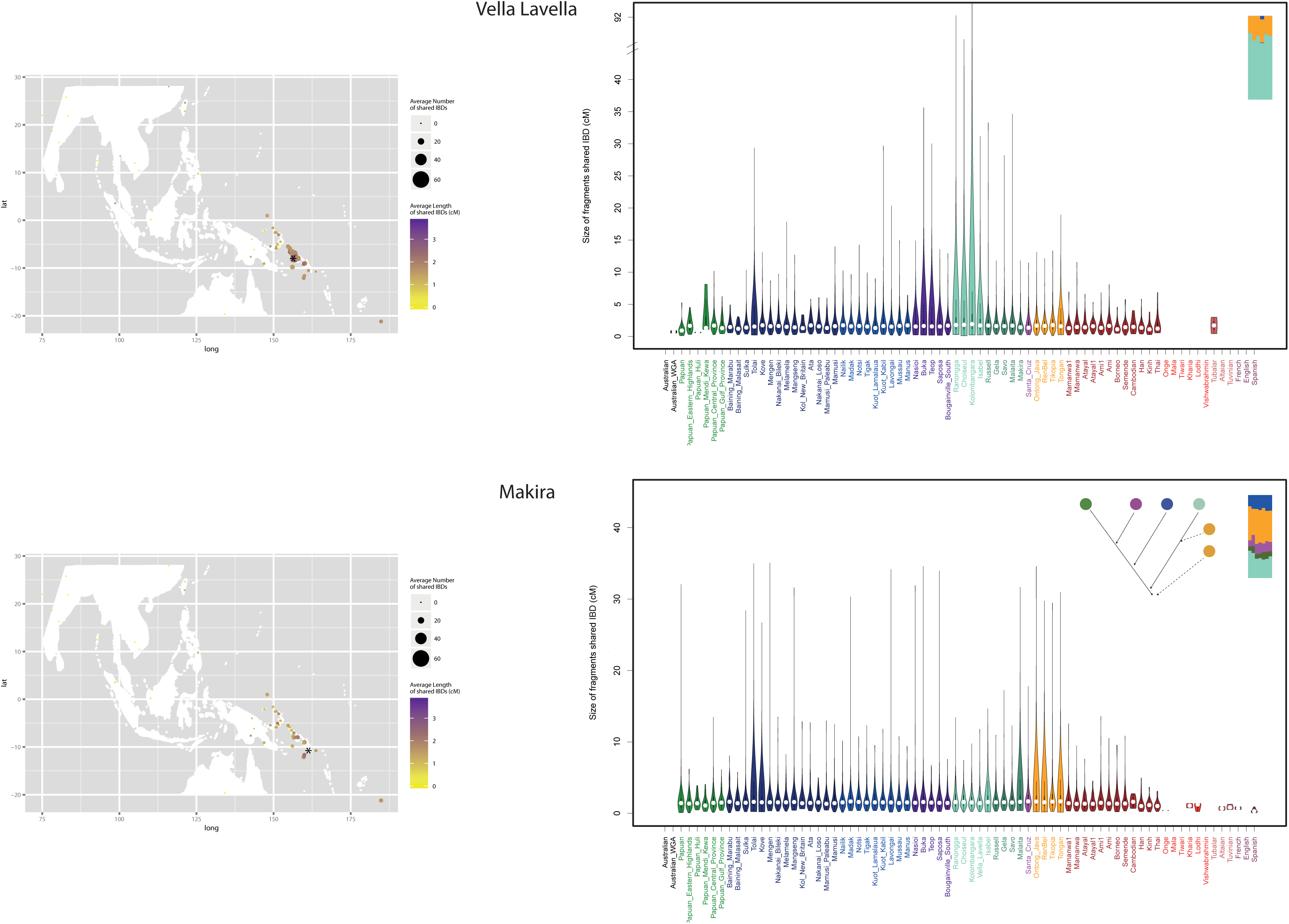
Dissimilarity in the pattern of sharing of IBD blocks between populations from western (A) and eastern (B) Solomons. Each data point represents the results for the comparison of the population marked with an asterisk (A: Vella Lavella; B: Makira) to each of the other populations in the dataset. Data points are placed on the map according to the sampling location of each population. The size of each circle is proportional to the mean number of IBD blocks shared, and the color intensity indicates the mean length of such shared blocks.

### Chronology of admixture events

Next, to better understand the chronology of gene flow across Near Oceania, we applied the Admixture History Graph (AHG) approach (Pugach et al. 2016),which elucidates the sequence of admixture events in populations with more than two ancestry components inferred by the ADMIXTURE analysis. The AHG approach is based on differences in the covariance between the various ancestry components inferred by ADMIXTURE (Pugach et al. 2016). Since the results of the ADMIXTURE analysis are sometimes ambiguous and do not necessarily reflect admixture (as discussed above), it is important to note that in the absence of gene flow the results of the AHG test are simply inconclusive, as no difference in covariance between the ancestry components is observable. Therefore, the AHG approach will not result in a false interpretation of the ADMIXTURE analysis.

As already discussed, most of the components inferred by the ADMIXTURE analysis in Oceania do not reflect distinct ancestries, yet an Austronesian-related signal is clearly present, as seen in the f3 results that indicate admixture involving Taiwan aboriginals (Table.S2). This signal is associated with the orange ancestry, and is seen in the highest frequency in PO (Fig.2). It therefore seems reasonable to interpret any AHG configuration in which the orange ancestry is not the most recent as suggesting additional post-Austronesian gene flow. In New Britain, distinguished by the higher frequency of the blue and pink ancestries, the AHG analysis infers Polynesian gene flow as appearing last only in the Mengen, Melamela and Mangseng groups (Fig.S13A). For the other populations, the AHGs are not completely resolved for the four ancestries, but suggest either the pink (present at high frequency in the Mamusi groups) or the aqua (present at high frequency in Bougainville) ancestry as the most recent. The results of the AHG analysis for populations from New Ireland and Lavongai are inconclusive, which corroborates evidence from the IBD-sharing analysis of continuous admixture and/or a complex history of gene flow for these populations. Notably, neither the AHG nor the IBD-sharing analysis suggest that the Polynesian ancestry is the most recent in these groups (Fig.S13B). All Bougainville populations with more than two ancestries inferred (Buka, Saposa and Teop) have identical AHG configurations, which infers the aqua ancestry as most recent (Fig.S13D), indicating substantial recent local gene flow. AHG analysis for the western SI populations was not possible, because they exhibit only two ancestry components. The eastern SI populations (Russell, Gela, Savo, Malaita and Makira) show more Papuan-related ancestry, and the AHGs were different than those inferred for the Bougainville populations, as the orange ancestry was coupled with the aqua (Bougainville) ancestry (Fig.S13D). For Makira the AHG is similar, but there is additional recent gene flow from an orange ancestry source, which is also corroborated by their sharing long IBD segments not only with Tonga (as is the case for the other islands in the eastern SI), but with the Polynesian Outliers as well (Fig.S13D). Finally, the AHG for Santa Cruz looks similar to those inferred for New Britain, but with additional gene flow from some Papuan-related source (Fig.S13E).

To summarize, the analyses based on the number and length of segments shared IBD and the AHG analyses strongly suggest that, with a few exceptions, most populations in Near Oceania exhibit a signal of post-Austronesian gene flow of diverse sources, both eastward from the BA and westward from Polynesia.

### Dating admixture events

To infer the date when (presumably) Austronesian-related ancestry entered Near Oceania, we first used PCAdmix (Brisbin et al. 2012) to infer local ancestry along individual chromosomes for all populations in the BA, SI and for Tongans. For this inference highlanders of PNG and either Taiwanese Aboriginals (as suggested by f3 and TreeMix analyses) or Polynesian Outliers (as suggested by ADMIXTURE) were used as the two source populations for the admixture. Once the ancestry blocks contributed by the two source populations were identified, we estimated the width of the ancestry blocks via the wavelet transform (WT) analysis (Pugach et al. 2011), and the date of admixture was inferred based on the simulations described previously (Pugach et al. 2011). When Taiwanese aboriginals are used as the source of Asian ancestry, the date of admixture inferred for all of the admixed populations falls between 77 and 104 (95% CI: 66 ‐152) generations ago, which using 30 years for the generation time (Fenner 2005) corresponds to ~ 2,300-3,100 years since admixture. These dates, which due to subsequent gene flow from Polynesians are likely more recent than the dates of initial admixture, are still within the chronological bounds of the Lapita period (Bellwood 2004) or just below them (Fig.6). Depending on the population, using Polynesians as the source of admixture results in admixture dates that are 150-1000 years younger, which is not surprising given that the Polynesians are themselves of dual Asian-Papuan ancestry (e.g. Kayser et al. 2008; Friedlaender et al. 2008; Wollstein et al. 2010). The admixture date is substantially more recent for the Tongans (~1500 years ago) because Tongans are themselves Polynesian, and there is very little differentiation between them and the Polynesian Outliers used as the parental group. With Taiwanese Aboriginals used as the parental group, the inferred date of admixture for Tongans is 83 (95% CI: 66-112) generations ago or ca. 2,500 years ago. This date of admixture is consistent with the dates of 90 generations (95% CI: 77-131) (Pugach et al. 2011) and 99 generations (95%CI: 19-267) (Wollstein et al. 2010) estimated previously by two independent methods for a diverse sample of Polynesian islands which included Tonga (Wollstein et al. 2010; Pugach et al. 2011). However, this date is older than an estimate of 50-80 generations ago obtained with the ALDER method (Loh et al. 2013) on the same Tongan samples (Skoglund et al. 2016). Similarly, all the admixture dates inferred using ALDER for the populations of Near Oceania appear to be 10-20 generations younger (Skoglund et al. 2016) than those inferred here using the WT method.

**Fig.6.**
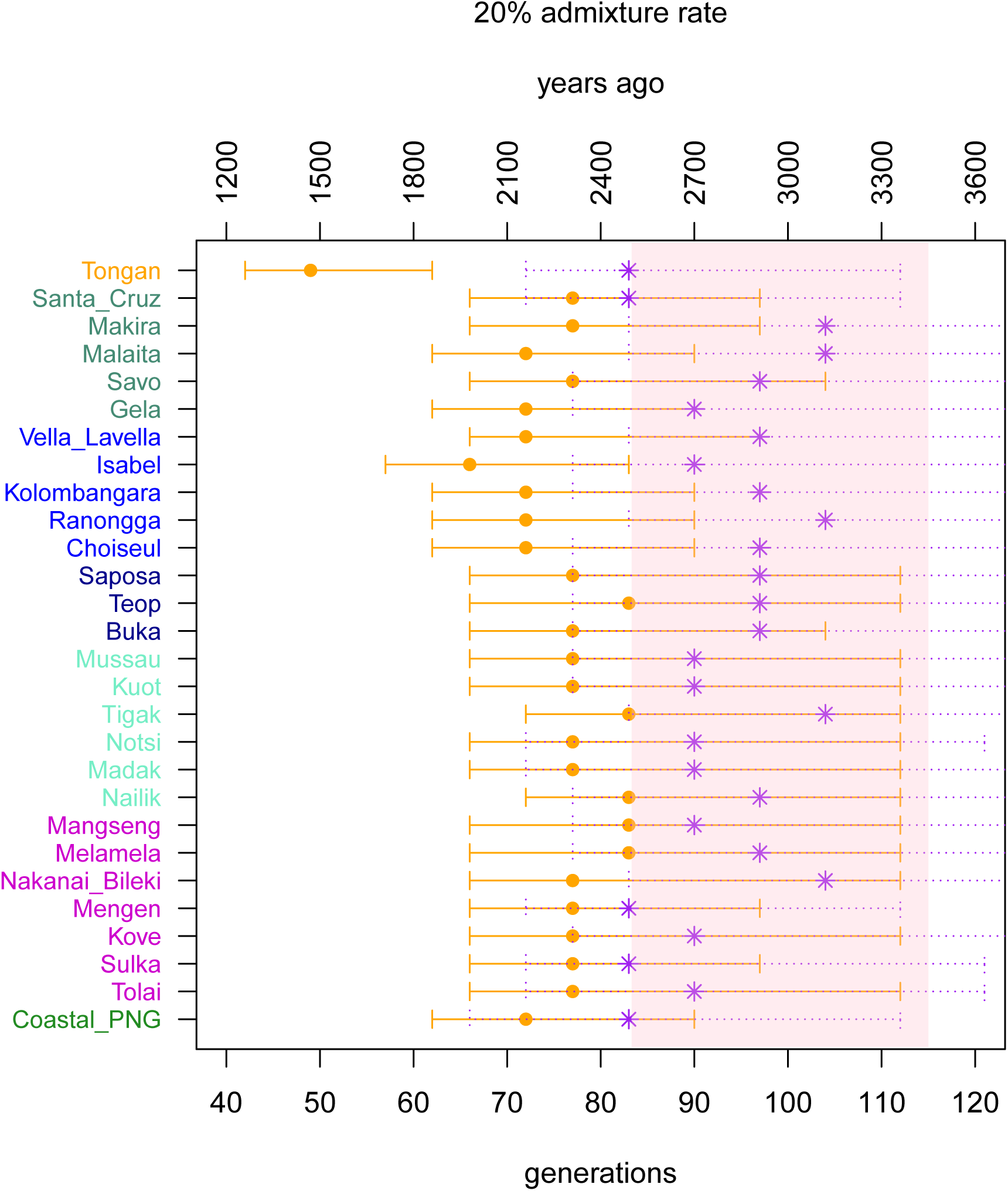
The dates of admixture inferred via the Wavelet Transform Analysis. In assigning local ancestry along individual chromosomes, we used the PNG Highlanders as a proxy for the Near Oceanian ancestry, while for the Austronesian ancestry we either used the Taiwan aboriginals (the inferred admixture dates are shown in purple) or the Polynesian Outliers (the inferred dates are shown in orange). Error bars represent the 95% confidence interval. The times of admixture are expressed in either generations or years, assuming a generation time of 30 years. The chronological bounds of the Lapita period are shown as a pink band.

To understand the source of this discrepancy we compared the WT method to ALDER using data simulated under a variety of different admixture scenarios. For this comparison, we used coalescent simulations implemented in the software MaCS (Chen et al. 2009). The basic simulation setup was as described previously (Pickrell et al. 2014 and Fig.S14). Briefly, for each admixture scenario we simulated data from nine populations, each consisting of 30 individuals with 10 independent chromosomes of 200Mb. One of the simulated populations was the product of an admixture between two other populations in the simulated dataset, with the admixture times set to 40, 70, 90, 100, 110, 120, 130, 140, 150 or 165 generations ago. Each simulated history was generated ten times. We then applied ALDER and the WT method to compare how each performs in recovering the simulated dates. Our results indicate that given a single episode of admixture, both methods perform reasonably well in recovering the simulated date (Fig.7). We then proceeded with the same simulation setup, but simulating two independent pulses of admixture from the same source population occurring at different time points. The first pulse of gene flow was set to have occurred 100, 120 or 140 generations ago, followed by the second pulse at 40, 60 or 90 generations ago, thereby simulating nine different admixture histories. As before, each of the admixture histories was simulated ten times. We then proceeded with ALDER and WT analysis to infer admixture dates. Note that while both methods have extensions which enable their use in situations involving multiple events of admixture from different sources (Pickrell et al. 2014; Pugach et al. 2016), here we are simulating multiple pulses of gene flow from the same source population. Our expectation is that a single estimate of the time of admixture would be inferred, and it would be a composite date reflecting both the older date of admixture as well as the more recent additional gene flow. The simulation results (Fig.7) indicate that in such situations ALDER tends to recover a date which is close to the most recent date of admixture, while the date estimated with the WT method is closer to the mid-point between the two events. It therefore seems likely that the more recent dates of Austronesian gene flow inferred by ALDER vs the WT method for Near Oceania and Tonga reflect the fact that after the admixture resulting from the initial Austronesian expansion, most of these populations experienced additional gene flow from a population(s) related to Austronesians, i.e. from Polynesia. This post-Austronesian gene flow is indeed strongly suggested by our analyses, in particular by the amount and length of IBD segments shared between Near Oceania and Polynesian Outliers and Tongans (Fig.S13).

**Fig.7.**
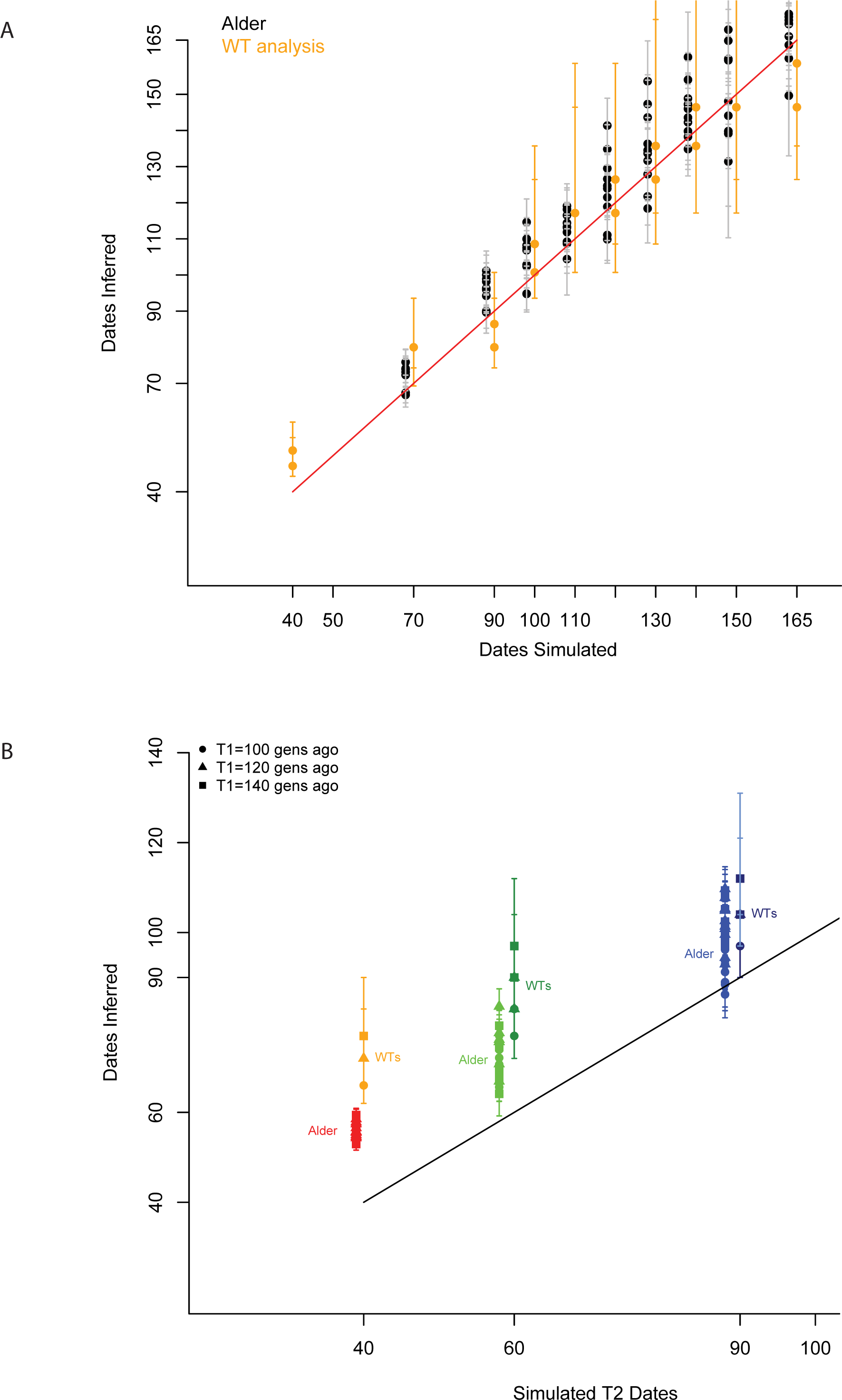
Performance of the ALDER and the Wavelet Transform method in recovering simulated dates of admixture. Both methods were applied to (A) ten simulated datasets with different admixture histories, but involving only a single instantaneous event of gene flow occurring 40-165 generations ago, (B) nine simulated datasets, generated using two pulses of gene flow from the same source, the more recent episode occurring 40, 60 or 90 generations ago, with the earlier episode of admixture 100, 120 or 140 generations ago. For both (A) and (B) each admixture time point was simulated ten times. The error bars denote the 95% confidence interval. While running both methods a single event of admixture was assumed. The estimates from ALDER and the WT method are slightly off-set from each other for better visibility of the results.

### Unusual genetic profile of the Santa Cruz islands and evidence for the “leapfrog” colonization from the Bismarcks

Although politically part of the Solomon Islands, the Santa Cruz islands lie beyond the easternmost part of the SI chain, and are separated from the other islands by 400 km of open ocean. Santa Cruz is thus one of the westernmost islands of Remote Oceania, which was only colonized when the Lapita people introduced navigational skills and technology which made long-distance ocean voyaging possible. As proto-Polynesians crossed into Remote Oceania, the Santa Cruz islands were probably the first landmass to have been colonized, with the archaeological sites suggesting human occupation at approximately 3.2 kya (Gross 2016). The languages of Santa Cruz belong to a primary branch of Oceanic languages (Næss 2006; Ross & Næss 2007; Næss & Boerger 2008) and are so distantly related to other Austronesian languages in the Solomons, that they were originally classified as Papuan with perhaps some Austronesian substrate (Wurm et al. 1978). From a genetic perspective, the origins of the Santa Cruz islanders remain a mystery, as they are remarkably different from all other Polynesian populations, harboring a high proportion of Near Oceanian mtDNA (Friedlaender et al. 2002; Duggan et al. 2014) and Y-chromosome (Delfin et al. 2012) haplogroups. Moreover, the high frequency of Near Oceanian mtDNA and Y-chromosome lineages cannot be explained by a recent population bottleneck or founder event, as there is great diversity within the Near Oceanian mtDNA and Y-chromosome lineages (Delfin et al. 2012; Duggan et al. 2014).

The unique genetic profile of Santa Cruz is also confirmed by the genome-wide data. As previously mentioned, the placement of Santa Cruz in the PC analysis is rather unexpected; given the geographic location, we would expect Santa Cruz to fall closest to the other populations of Remote Oceania. Instead they are positioned closest to populations from New Britain in the BA, and remain so positioned at higher PCs (Fig.S3). The results of the ADMIXTURE analysis are equally perplexing - Santa Cruz is the furthest of the Solomon Islands from PNG, yet reveals the highest frequency of Papuan-like ancestry components, with only 5% coming from Polynesia (Fig.2).

The sharing of IBD segments within Santa Cruz is low, especially in comparison to the Polynesian Outliers (Fig.S15), which is further evidence against recent population bottlenecks (Gusev et al. 2012). The absence of a population bottleneck in Santa Cruz is also evident from the pattern of genome-wide LD and ROHs; namely there is no evidence of elevated LD or of excess of ROHs as observed in Baining and some of the Polynesian Outliers (Figs.S16 and S12), and as is typically associated with temporary reductions in population size (Pritchard & Przeworski 2001). Furthermore, it has been shown previously that the slow rate of decay in abundance of small IBD segments (2-10 cM) in comparison to longer segments is indicative of founder events (Palamara et al. 2012); such slow decay is observed in Polynesian Outliers, but not in Santa Cruz, similarly confirming that in addition to the absence of a population bottleneck, there is no evidence to support a founder event in Santa Cruz either.

Next, we examined the decay in correlation of genome-wide LD between Santa Cruz and other populations. In general, the patterns of correlation in genome-wide LD between populations can be used to estimate their time of divergence (Sved et al. 2008; McEvoy et al. 2011). These estimates however are known to be heavily biased downwards (making the times of divergence too recent) by substantial levels of gene flow between populations following their separation, and by drift in smaller populations (Sved et al. 2008); both issues are highly relevant for Near Oceanian and Polynesian populations, as is evident from all our analyses. So instead of estimating the times of divergence in absolute terms, we have used this as a relative measure of how close the populations are to each other. The basic logic behind this analysis is that following divergence, the correlation in genome-wide LD between the two daughter populations will decay exponentially over time, with the rate of decay dependent only on the recombination distance between the markers (McEvoy et al. 2011). For this analysis, we have computed the correlation between the LD values for each pair of populations and for each recombination distance category, and then determined at which recombination distance the correlation decays to zero, surmising that for populations with recent genetic ties this correlation should persist for longer genomic distances (i.e. there would be less time for recombination to break down the correlation). Indeed, the correlation in LD between the French and any population from Oceania decays to zero at about the same recombination distance of ~ 0.4 cM, suggesting that the French are approximately equally related to all populations in the analysis (Fig.S17A). In contrast, the correlation in LD between two Polynesian outliers, Rennell & Bellona (RB) and Ontong Java, persists for much longer recombination distances, more than 0.8 cM, while the correlation between RB and populations from the BA and the SI decays much faster, becoming zero at a distance between 0.45-0.65 cM (Fig.S17B). Interestingly, this correlation persists for longer distances between Santa Cruz and the BA (0.57 cM with Sulka and Notsi), than for Santa Cruz and the SI (0.55 cM with Makira; 0.52 cM with Choiseul) (Fig.S17C). To compute the significance of this difference, i.e. if it is consistent with zero, we used a weighted block jackknife procedure (Busing et al. 1999) by sequentially dropping each of the 22 chromosomes (see Methods) and recomputing the correlation in LD between Santa Cruz and BA, and Santa Cruz and SI, for each of the runs. A jackknife estimate of the difference in the rate of the correlation decay is highly significant with Z = 3.96 and Z=6.12 standard errors from zero for the difference between SC and the BA vs SC and Makira, and between SC and the BA vs SC and Choiseul, respectively.

To summarize, in all analyses Santa Cruz falls consistently closer to populations from the Bismarcks than to those from the Solomon Islands, and appears to be quite different from the other populations of Polynesian origin. The high frequency of Papuan-like ancestry in Santa Cruz, and the gradient of this ancestry diminishing across the eastern SI in an east-west direction, is in accordance with previous suggestions (Sheppard 2011) that the first people to expand into Remote Oceania did so in a “leapfrog” manner from the Bismarcks, completely bypassing the SI. The SI were then subsequently settled in a bidirectional manner, underlying the clear distinction in the genetic background between western and eastern parts of the chain (Fig.5). Notably, this distinction that we observe based on genetic evidence is mirrored by a high-order linguistic break, known as the Tryon-Hackman line (Ross 1988; Sheppard & Walter 2006), which demarcates the boundary between the Western Oceanic grouping of Austronesian languages (northern New Britain and New Ireland, and the northern and western Solomons to Isabel), and the Central/Eastern Oceanic grouping, which includes all Austronesian languages spoken to the east of this line (Lynch et al. 2002). There are also cultural differences between SI populations on either side of the Tryon-Hackman line (Sheppard & Walter 2006); the genetic evidence provides strong support for bidirectional settlement of the SI, and thus a potential explanation for the Tryon-Hackman line.

In conclusion, our analyses support the view that the initial dispersal of people from Bismarck Archipelago into Remote Oceania occurred in a “leapfrog” manner, completely by-passing the main chain of the Solomon Islands, and that the colonization of SI proceeded then in a bi-directional manner, underlying the divergence between western and eastern Solomons, in agreement with the Tryon-Hackman line. Santa Cruz remains a puzzle, exhibiting strong ties to the Bismarcks and high levels of Papuan ancestry not found elsewhere in the SI, yet also not exhibiting any signal associated with a founder event. It appears there was substantial Papuan-related gene flow to Santa Cruz that bypassed the rest of the SI. We also report ubiquitous recent post-Austronesian gene flow across the Solomons, both from the west and from the east. Finally, as more and more studies of human genetic variation emerge based on dense geographical sampling, it becomes apparent that geographically-close populations can have quite different histories (e.g. in Siberia (Pugach et al. 2016) and Indonesia (Hudjashov et al. 2017)), so dense genome-wide data and fine scale sampling are needed to unravel all the complexities of human population history.

## MATERIALS AND METHODS

### Samples and Genotypes

The dataset includes samples from 50 populations from Near and Remote Oceania (Fig.S1): aboriginal Australians, 3 populations from the highlands and 3 populations from the coastal parts of mainland Papua New Guinea, 23 populations from the Bismarck Archipelago (islands of New Britain, New Ireland, Lavongai, Mussau and Manus), 5 populations from Bougainville, 11 populations from the Solomon Islands, 3 Polynesian “outlier” populations (Rennell and Bellona, Ontong Java, and Tikopia) sampled in the Solomon Islands, and Tonga. For comparative purposes we also included samples from Mainland and Island South East Asia, India and Western Eurasia (Table.S1 and Fig.S1). All samples were genotyped previously (Lazaridis et al. 2014; Qin & Stoneking 2015; Skoglund et al. 2016) on the Affymetrix Human Origins SNP Array, which includes 13 panels of SNPs ascertained in different populations, including Papuans (Patterson et al. 2012). Data curation was performed as described previously (Pugach et al. 2013). After quality filtering and data integration, the full dataset comprised 823 individuals and over 620,000 autosomal SNPs. Not all data were used for all of the analyses.

### Principal Components Analysis

PCA was performed using the StepPCO software (Pugach et al. 2011) on the entire dataset, on a subset excluding populations of Western Eurasia and India, and on a subset comprising only the Oceanian populations.

### ADMIXTURE

To infer individual ancestry components and admixture proportions we used the ADMIXTURE software (Alexander et al. 2009). The analysis was performed for the entire dataset and for the dataset comprising only the Oceanian populations. The LD pruning for the analyses was done using the PLINK tool (Purcell et al. 2007) with the following settings, which define window size, step and the r^2^ threshold: ‐indep-pairwise 200 25 0.4 (Rasmussen et al. 2011), which reduced the data set to 215,869 markers. We ran ADMIXTURE for K = 3 through K = 12 for the full dataset, and from K = 3 through K = 8 for the Oceanian subset. One hundred and twenty independent runs for each value of K were performed for each dataset. We confirmed consistency between runs and used the cross-validation procedure implemented in ADMIXTURE to find the best value of K (Fig.S6). In addition, to evaluate how the choice of SNPs affects the results, for the full dataset we performed ten independent runs for each value of K from K = 7 through K = 11 using the SNPs from each of the thirteen ascertainment panels (Patterson et al. 2012) separately. Results for each of the panels for K=10 are shown in Fig.S8.

### 3-population test

Formal tests of admixture were carried out using the f3 statistics. These f3 statistics were calculated with the qp3Pop script from the AdmixTools package (Patterson et al. 2012).

### Inferring Segments IBD

The data were phased using BEAGLE v3.3.2. Segments that are identical by descent (IBD) were inferred using the fastIBD method implemented in BEAGLE (Browning & Browning 2007; Browning & Browning 2011). We ran the algorithm 10 times with different random seeds. The results were then combined and postprocessed as described previously (Ralph & Coop 2013; Pugach et al. 2016) to extract only those IBD segments seen at least twice in the ten BEAGLE runs and which had a significance score lower than 10^−9^. We also merged any two segments separated by a gap shorter than at least one of the segments and no more than 5 cM long, thus removing artificial gaps potentially introduced because of low marker density or possible switch error during phasing (Ralph & Coop 2013; Pugach et al. 2016). All results were adjusted for differences in sample size, and blocks shorter than 2cM were excluded from some of the analyses (Ralph & Coop 2013). To run PC analysis based on the results of the IBD calculation, we used the ade4 R package (Dray & Dufour 2007); the inverse of the matrix with the number of IBD blocks shared by each pair of individuals was used as a distance matrix (Pugach et al. 2016).

### ROH

Runs of homozygosity (ROH; consecutive stretches of homozygous SNPs) provide information about relatedness among individuals and population isolation. Runs of homozygosity were identified using PLINK (Purcell et al. 2007) with default settings.

### Linkage Disequilibrium and Correlation in LD

Genome-wide linkage disequilibrium (LD) was estimated by binning the genome-wide data into 50 evenly spaced recombination distance categories (0.005–1.000 cM). For each population and for every pair of SNPs within each distance category, we calculated the squared correlation (rLD^2^) in allele frequencies by randomly selecting 7 individuals from each population (to ensure equal number of individuals per population, since the smallest sample size in this dataset was 7) and further adjusting the measurement for each pair of SNPs to account for missing data (McEvoy et al. 2011; Pugach et al. 2013). The correlation between the LD values was computed for each pair of populations and for each recombination distance category (McEvoy et al. 2011; Pugach et al. 2013). Standard errors for the difference in the distance over which the correlation between the LD values decays to zero were computed using the weighted jackknife approach (Busing et al. 1999) with chromosome-sized blocks and weighting each replicate by the total number of SNPs in each of the runs.

### Admixture History Graphs

For populations exhibiting evidence of multiple admixture events, the order of the admixture events was inferred using admixture history graphs as described previously (Pugach et al. 2016). All tests for covariance between the ancestry components were based on the results for K = 5 (Fig.S13) of the ADMIXTURE analysis based on the Oceanian populations only.

### Dates of admixture inference

To infer local ancestry along individual chromosomes and obtain the genome-wide block-like signal of admixture we used a PCA-based approach implemented in the PCAdmix software (Brisbin et al. 2012). In choosing the proxies which would best represent the parental populations we followed the results of the 3-population test and ADMIXTURE analyses. Specifically, we used the PNG highlanders to represent the Near Oceanian ancestry component, and either the Polynesian Outliers or the Taiwan Aboriginals as the proxy for the Austronesian ancestry component. After retrieving the block-like admixture signal using PCAdmix, which assigns ancestry along individual chromosomes to either of the two source populations, we applied wavelet-transform analysis and used the wavelet transform coefficients to infer time since admixture by comparing the results to simulations, as described previously (Pugach et al. 2011).

### Comparison between the Wavelet Transform and ALDER methods

We used simulated data to assess the performance of these two methods for dating admixture events. For each admixture scenario we simulated data from nine populations with the MaCS software (Chen et al. 2009), each consisting of 30 individuals with 10 independent chromosomes of 200Mb. One of the simulated populations was the product of an admixture between two other populations in the simulated dataset (Pickrell et al. 2014 and Fig.S14), with the admixture rate set to 0.2. For the one pulse admixture scenarios, the admixture times were set to 40, 70, 90, 100, 110, 120, 130, 140, 150 or 165 generations ago, thus generating ten distinct admixture histories. For the scenarios involving two pulses of gene flow from the same source, the earlier event was set to 100,120 or 140 generations ago, and on each of these backgrounds an additional admixture event was set to occur 40, 60 or 90 generations ago, thus producing nine distinct admixture histories. Each simulated admixture history was generated ten times. As an example, below is the MaCS command line used to simulate two pulses of admixture from the same source population, occurring 90 and 140 generations ago:
macs 540 200000000 ‐t 0.00004 ‐r 0.0004 ‐I 9 60 60 60 60 60 60 60 60 60 0 ‐em 0.0035 1 7 8000 ‐em 0.003525 1 7 0 ‐em 0.00225 1 7 8000 ‐em 0.002275 1 7 0 ‐ej 0.0125 7 8 ‐ej 0.0125125 1 2 ‐ej 0.0125125125 4 5 ‐en 0.0249 8 0.02 ‐ej 0.025 8 9 ‐en 0.0249249 2 0.02 ‐ej 0.02525 2 3 ‐en 0.0249249249 5 0.02 ‐ej 0.0252525 5 6 ‐en 0.0748 9 0.01 ‐ej 0.075 6 9 ‐en 0.1498 9 0.01 ‐ej 0.15 3 9

When inferring the times of admixture based on the wavelet transform analysis, PCAdmix (Brisbin et al. 2012) was first applied to the simulated dataset to determine local ancestry along the chromosomes, with the source populations defined by the simulation set-up (Fig.S14). The WT analysis was then performed as described previously (Pugach et al. 2011), with wt coefficients compared to those obtained with simulations (Pugach et al. 2011) based on either the admixture rate of 0.2 for the one-pulse or 0.4 for the two-pulse scenarios. ALDER was run with default parameters, with all “unadmixed” simulated populations supplied as potential sources.

### DATA AVAILABILITY

To comply with the informed consent under which the samples were obtained, we make the data available upon request by asking the person requesting the data to agree in writing to the following restrictions: 1) The data will only be used for studies of population history; 2) the data will not be used for medical or disease-related studies, or for studies of natural selection; 3) the data will not be distributed to anyone else; 4) the data will not be used for any commercial purposes; and 5) no attempt will be made to identify any of the sample donors.

## ACKNOWLEDGMENTS

We thank all individuals who contributed samples for this study. We thank David Reich for assistance with genotyping, and Murray Cox and Cosimo Posth for valuable discussion. This research was funded by the Max Planck Society.

## AUTHOR CONTRIBUTIONS

IP and MS conceived and designed the experiments. DAM, FRF, and JSF provided reagents. IP performed the experiments, analyzed the data with assistance from ATD, and contributed analytical tools. IP and MS wrote the paper with input from all authors.

